# How Efficient is the k-means Clustering to Analyze the CT images of Pyogenic and Amoebic Liver Abscess?

**DOI:** 10.1101/2022.08.06.503068

**Authors:** Subhagata Chattopadhyay

**Affiliations:** Independent researcher

**Keywords:** Amoebic liver abscess, Pyogenic liver abscess, Image processing, Teleradiology, k-means clustering

## Abstract

Liver abscesses are well-delineated pus-filled lesions. Two common types are Amoebic liver abscesses (ALA), caused by protozoa called entamoeba histolytica while several pus-forming bacteria cause pyogenic liver abscesses (PLA). Both cause debilitating morbidities and are diagnosed by pus culture-sensitivity tests. A contrast CT abdomen shows well-demarcated lesions in the liver. The telemedicine practice is on the rise where image processing is becoming a part and parcel of teleradiology to fill the gap between the number of radiologists versus the large patient pool. Cluster-based image segmentation is a useful step in grouping the image into the desired number of clusters. The k-means clustering (k-MC) technique is one popular method, used in this study on ALA and PLA contrast CT images. it observes that with the desired 2-clusters – a) normal liver tissue and b) the pus-filled tissue) parameters, the algorithm gives better results in PLA.

## Introduction

With an incidence rate of 2-3/100,000 population liver abscess is commoner in males between 40-60 years of age and the portal vein is the key porter of the infection, mostly to the right lobe of the liver, close to the diaphragmatic dome [1]. The two most common types of liver abscesses are amoebic and pyogenic liver abscesses, in short, ALA and PLA, caused by *entamoeba histolytica* and *escherichia coli*, respectively although other pyogenic bacteria are also held responsible [1]. Abdominal contrast computerized tomography (CT) provides a high-quality image of liver abscesses with a diagnostic accuracy of 96% compared to conventional abdominal ultrasound having a diagnostic accuracy of around 76% [2].

Teleradiology is becoming a part and parcel of telemedicine to bridge the gap between the small number of radiologists versus a large number of populations in need. Its advantages are manifold when accessing healthcare from distance is the key issue that is commonly noticed in a geographically vast country like India. Within teleradiology, again, a radiologist has to handle a high volume of images. The image processing technique here comes into play as a method of not only increasing the processing speed but also identifying the region of interest (ROI) in the image [3]. In many places, healthcare providers do not have any radiology exposure. To them, automatic ROI detections are helpful to decide on the referral of a case to a higher facility without much delay [3].

Segmentation methods are useful for medical image processing to find out the ROIs automatically. Among many segmentation techniques, k-means clustering (k-MC) is a popular one. It randomly assigns the ‘centroid-pixels’, iteratively, and then measures the Euclidean distances of the remaining pixels from the centroid till no more pixel remains unclustered. It is user-friendly as it can define the number of desired clusters and is quite fast to converge [4].

### k-MC algorithm

It is an unsupervised learning algorithm when the classes/clusters are unlabelled, such as in this case.

Steps-

#### Start

1. assign the number of ‘k’ clusters
2. select random cluster centers (centroids) as ‘k’ points iteratively
3. measure the distance of each data point (here it is a pixel) from the centroid iteratively and accommodate if the distance is less than a cut-off value
4. accommodate all data points into ‘k’ clusters iteratively till no data point is left out

#### Finish

The *objective* of this study is to (a) check how efficient is k-MC in segmenting the lesions in the ALA and PLA CT images from the liver tissue and (b) the pathological correlations.

### Image processing

The ALA and PLA contrast CT images (.jpg) is acquired from [5] and the published research paper by [6], respectively. The images are then processed by grayscaling followed by the local thresholding, edge-detection with Sobel horizontal and vertical methods, and finally clustering-based segmentation with the k-MC method. Before clustering, descriptive statistics (minimum value, maximum value, mean value, median value, value of standard deviations, and quartile values) of the pixel data for each image have been calculated. Histogram plots (pixel value on the x-axis vs frequency of occurrence on the y-axis) of each image are also conducted to note the pixel density (size and its respective frequency). All programming is accomplished using *python 3*.*8*.*3* with the pre-loaded image processing packages, which run on Windows 10 pro OS platform. The outputs can be seen comprehensively in Fig. 1 (ALA image processing) and 2 (PLA image processing), respectively. It is important to note that black zones represent the minimum pixel values, i.e., zero while the brighter zones have higher values close to 255.

**Fig. 1.**
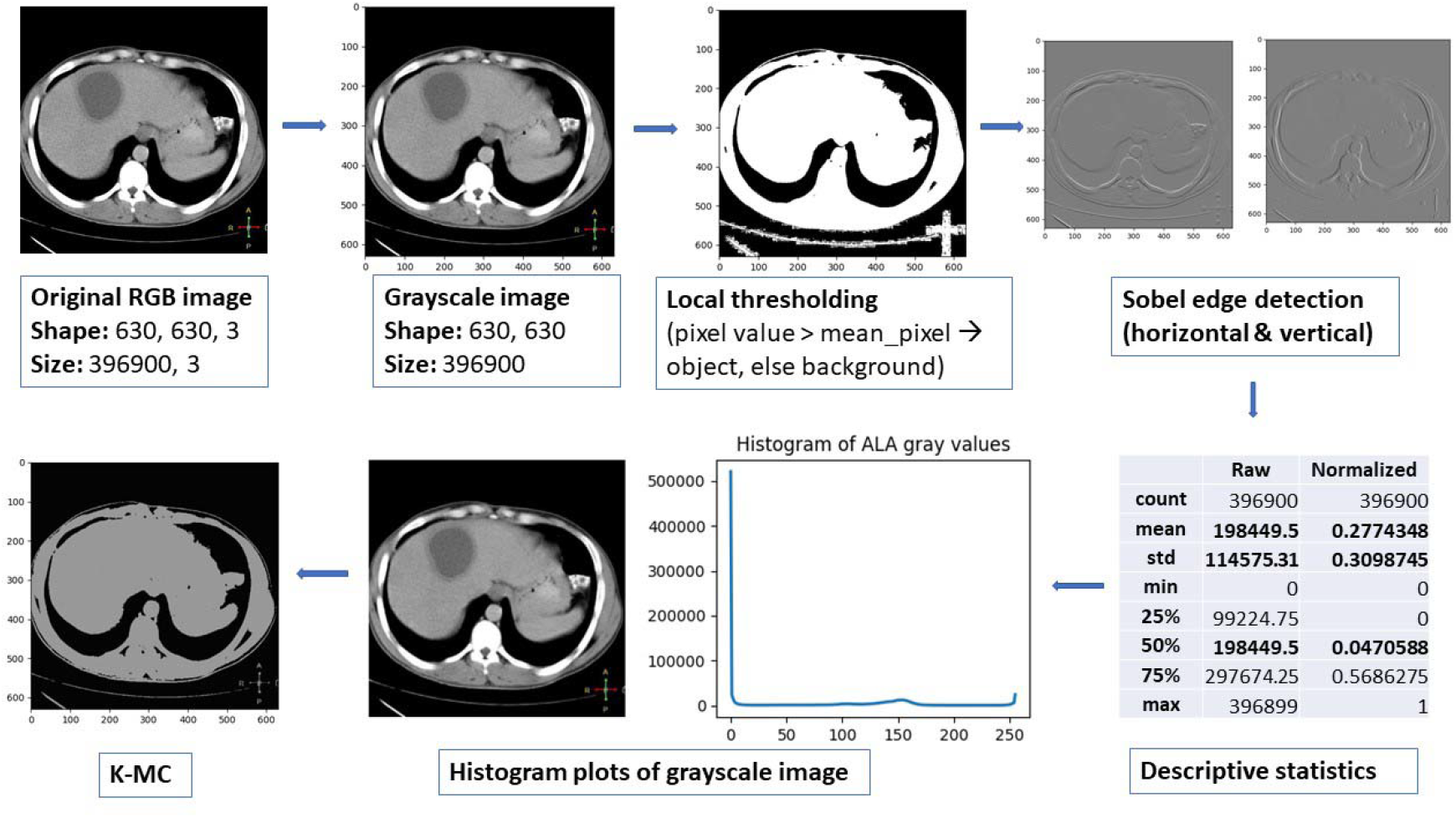
Image processing till the k-MC in ALA contrast CT image.

Fig. 1 shows a comprehensive picture of how the CT image of ALA has been processed till k-MC clustering. The lesion can not be detected in any of the processes such as the local thresholding, Sobel edge detection, and k-MC. The histogram plot is a dull line that corroborates the fact that there is no difference in the tone between the lesion and the liver parenchyma. The normalized mean and median pixel values are 0.277 and 0.04 with a standard deviation of 0.309.

Fig. 2 shows a comprehensive picture of how the CT image of PLA has been processed until k-MC clustering (2-clusters) is done in the event of ALA. The lesion can not be detected only by the local thresholding. Other more sophisticated techniques, e.g., Sobel edge detection, and k-MC could detect the lesions effectively. The histogram plot is not a dull line as the ALA histogram plot. It shows tiny spikes between 0 and 50 marks which corroborates the fact that there is an obvious difference in the tone between the pus-filled lesions (darker region) and the liver parenchyma (relatively brighter region).

**Fig. 2.**
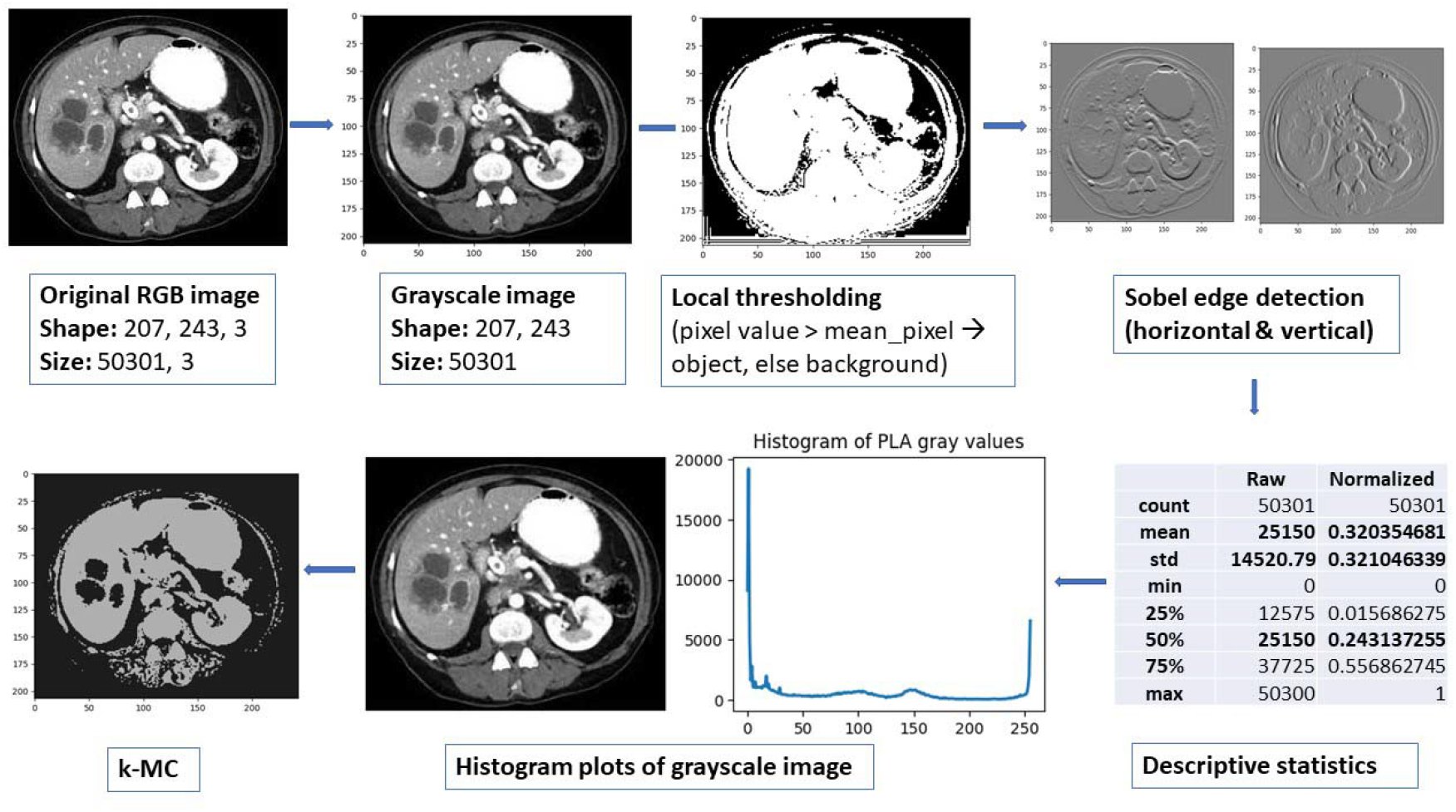
Image processing till the k-MC in PLA contrast CT image.

### Pathological correlation of k-MC findings

In the case of ALA, the pus is typical of a chocolate brown color called as ‘anchovy sauce/paste’ type, i.e., much thicker in consistency. The pus consists mostly of dead and deformed liver cells (hepatocytes) and trophozoites of entamoeba on the wall of the abscess. There are a few red blood cells (RBC) and leucocytes or white blood cells (WBC) that can also be found [7]. It is important to state that superadded bacterial infection is much rare in ALA [8]. On the other hand, pus in PLA is due to overwhelming bacterial infection and hence it contains a high amount of dead WBCs (especially the neutrophils), a few necrosed hepatocytes, and an abundant number of RBCs. Hence, the pus is purulent like sputum but not as thick as the anchovy paste seen in ALA [9].

From this experiment, it can be ascertained that a high amount of blood cells in PLA has probably played a crucial role in manipulating the pixel values of the abscess images helping the k-MC method to successfully cluster the abscesses from the liver parenchyma. In the case of ALA, the mean pixel value is very high which is 198449.5 (i.e., more inclined towards the brighter pixel side) compared to PLA which has a mean pixel value of 25150, which is 1/8^th^. times the former i.e., more inclined towards the darker pixel side. Hence, in PLA, the k-MC algorithm can produce a sharp contrast between the pus-filled abscess (darker pixels) and the liver tissue (relatively brighter pixels), but not in ALA as the pixel values in the abscess are almost similar to the liver tissue.

### Conclusions and future work

The paper aims to test the k-MC algorithm to differentiate between the ALA and PLA contrast CT images and presents its preliminary results. It has also focused to derive the correlations between the k-MC result and the microscopical nature of the pus, especially the components in it. Such type of study has not yet been reported to the best of the knowledge of the author. The study observes that k-MC is a suitable algorithm to differentiate the image of any pyogenic lesions where WBCs and RBCs are plentiful. On the other hand, the pus in ALA is principally composed of necrosed hepatocytes and trophozoites giving a homogeneous texture like the liver tissue. This is an interesting finding and could be useful for image processing in teleradiology practice to assist rural populations in developing nations where ALA and PLA cases are not infrequent. However, such observation requires further testing and validation on a large set of ALA and PLA images.

## Abbreviations

ALA: Amoebic liver abscess
CT: Computerized tomography
k-MC: k-Means clustering
PLA: Pyogenic liver abscess
RBC: Red Blood Cells
WBC: White blood cells

